# Steeper or Faster? Impacts of Similar-intensity Running Conditions on Heart Rate Variability

**DOI:** 10.1101/2024.02.23.581819

**Authors:** Vivian MCA ElDash, Ingrid Machado Cusin Ahmed ElDash, Fernando Silveira Marques, Jose E S Natali, Jose Guilherme Chaui-Berlinck

## Abstract

Increases in speed or slope promote different adjustments in the locomotor system and lead to a higher heart rate (HR). Because heart rate variability (HRV) is known to respond to exercise intensity and to biomechanical stimuli, we aimed to answer whether HRV would be sensitive to changes in speed and/or slope under similar HR. We hypothesize that HRV would depend solely on HR. To test this hypothesis, 6 healthy male runners (age 30 ± 7.5 yrs.) were recruited and a velocity V, with similar HR/HR_max_ was selected per volunteer. Changes in HR were referred to these particular velocities V, resulting in comparable exercise intensities. HRV was estimated through standard deviation (sdt) and root mean square of successive RR intervals (rmst), normalized per volunteer. Four running conditions were compared. We found that rmst decreased with speed and slope, with significant interaction between the independent variables. The run {0.7 V, 0% slope} was different from the remaining 3 conditions, which, in turn, were no different from one another. Thus, this estimator did not show a distinct sensitivity to variations in speed or slope. On the other hand, sdt decreased with both speed and slope, with no significant interactions. The runs performed at {0.7 V, 0% slope} and {0.7 V, 6% slope} were different from those at {1 V, 0% slope} and {1 V, 6% slope}. Therefore, sdt presented different values under similar HR. This finding leads us to conclude that cardiac control operates sensitively due to the mode of the changes imposed on the metabolic demand.

**Summary Statement:** Does heart rate variability respond similarly to running in different combinations of speed and slope? While one estimator depended on the exercise intensity, another was sensitive to the running conditions.

## INTRODUCTION

The capacity to travel long distances, particularly by running, has been accepted as a key component in the evolution of the genus *Homo*, with evidence of endurance running appearing in the fossil record as early as 2.6 million years ago (Bramble & Lieberman, 2004). The selective pressure behind the evolution of endurance running is still controversial (Pickering & Bunn, 2007), however it has been hypothesized that it allowed early hominids to have better access to protein sources, either through hunting or scavenging (Bramble & Lieberman, 2004). Paleoenvironmental reconstructions indicate that endurance running evolved in the savanna-woodlands, a mosaic ecosystem composed of closed forest-like patches and open grassland areas. Also, most of the African savanna is composed of flat plains having isolated hills (inselbergs) scattered in the landscape.

From that initial terrain, migration would lead to many diverse geographic scenarios, and locomotion would include transitions from level ground to slopes and vice-versa. Therefore, regardless of which was the selective pressure, endurance running in *Homo* evolved in a diverse environment and early hominids most likely ran in a variety of conditions, which must represent different physiological stimuli.

Running on a positive slope is associated with an increase in total muscle volume activated in the lower extremity (Sloniger et al., 1997), with a tendency to mid/forefoot strike, higher stride frequency and decreased stride length when compared to level running, resulting in an increased internal mechanical work, shorter aerial phase duration, longer time spent in stance (Padulo et al., 2013; Vernillo et al., 2017). Up to around 90% maximum velocity, speed increments are mainly met by increased stride length, resulting in more time spent on the swing phase (Dugan & Bhat, 2005; Mann & Hagy, 1980; Novacheck, 1998; Stöggl & Wunsch, 2016). Increased stride frequency will be more significant in attaining higher speeds only when velocity approaches maximum. Under such circumstances, total gait time is reduced.

If, on the one hand, running faster or on a steeper surface causes notably different responses in the musculoskeletal system, on the other hand, those stimuli are both associated with increased metabolic demand at the organismal level. When running uphill, additional energy is required to elevate the center of mass and to compensate for a reduction in the maximum possible elastic energy stored and released by tendons (Vernillo et al., 2017). For example, Cavagna & Kaneko (1977) and Minetti et al. (2002) found that the oxygen consumption rate increased linearly with speed. Minetti et al. (2002) also reported energy expenditure per meter to change according to the slope, but not irrespective of speed. Moreover, Padulo et al. (2013) observed an increase in heart rate (HR) by 5% (from 148 BPM to 155 BPM) and 15% (reaching 170 BPM) when running at 2% and 7% incline, respectively, compared to level.

Under primarily oxidative metabolism and steady-state conditions, the heartrate (HR) is a good linear indicator of metabolic rate (Green, 2011; McArdle et al., 2007). Considering that changes in both speed and slope have a pronounced impact on energy expenditure, it comes as no surprise that several different combinations between these two variables can result in similar metabolic demands, and this might be indicated by similar heart rates. In this sense, the study of Jones & Doust (1996) reports some interesting experimental findings. Nine trained male runners engaged in the study, and the experimental protocol consisted of a series of 6-minute runs on a treadmill at 6 velocities (2.92, 3.33, 3.75, 4.17, 4.58, and 5.0 m/s). The procedure was repeated on different days for 0%, 1%, 2%, and 3% incline. They observed, for instance, that running at 3.75 m/s at 3% grade leads to the same HR as 4.17 m/s at 1% grade (153 BPM). Also, a 3.33 m/s speed at 0% grade (124 BPM) was similar to 2.92 m/s at 2% grade (126 BPM).

HR is modulated by the autonomic nervous system (ANS), both the sympathetic and parasympathetic branches innervate the heart and the autonomic control responds to a diversity of afferences, including baroreceptor, chemoreceptor, humoral, muscular, and environmental inputs (Dong, 2016).

Even under steady-state conditions, the time interval between successive heartbeats is not constant. The pattern associated with heart rate variability (HRV) is typically nonlinear, nonstationary, multiscale variable, and temporarily asymmetric, making the time series of the HR a remarkably complex signal. Such complexity is thought to result from HR control mechanism response to internal and external stimuli. A periodic or random signal indicates malfunctioning of control mechanisms and/or of the controlled system (the heart itself), as in pathological conditions (Goldberger et al., 2002). Therefore, how HRV behaves under different conditions can shed light on cardiac control and the operation of the cardiovascular system (Dong, 2016; GUIDELINES, 1996; Perini & Veicsteinas, 2003).

The interpretation of HRV under exercise is still controversial (Bailon et al., 2013). HRV is inversely proportional to HR (Bailon et al., 2013; Michael et al., 2017) and exercise intensity seems to be the most important factor determining HRV in this context (Michael et al., 2017). Nevertheless, at high-intensity exercise, components associated with the physical activity itself, such as cadence and ventilatory frequency, influence the HRV and should be taken into consideration when interpreting data (Alikhani et al., 2017). For example, results from graded cycling tests in different fitness groups indicate that, under low-intensity exercise, the component associated with respiratory sinus arrhythmia (RSA) is the most important contributor to total power in the frequency domain (Bailon et al., 2013; Blain et al., 2009) something numerically equivalent to the variance in the time domain (see Parseval’s relation, Oppenheim et al., 1996). As for maximal graded exercise, around 40% of HRV was associated with cadence, and 60% with RSA (Blain et al., 2009).

Considering the importance of both exercise intensity and biomechanics in the modulation of HRV and how they relate to running, it would be relevant to analyze the influence that changes in speed and slope induce in HRV, particularly under the same HR.

### Objectives and hypotheses

In this study, we aimed to answer the following questions: How does HRV respond to increments in speed and/or slope during running? What is the magnitude of those responses? Is HRV more sensitive to a given factor (speed or slope)?

On a physiological basis, we hypothesize that HRV will decrease as speed and/or slope increases and proportionally related only to the heart rate.

## MATERIALS AND METHODS

### Subjects

Subjects were healthy males (n = 6), aged from 22 to 43 years old, see anthropometric data in Table 1. Criteria of inclusion: be engaged in running at least twice a week for a minimum of 1 year and log 25 or more kilometers per week. Criteria of exclusion: chronic diseases such as diabetes or hypertension or any disease of the cardiovascular and/or locomotor systems, including injuries in the past 6 months. A 12-lead clinical electrocardiogram was performed before the experimental protocol. All volunteers gave written informed consent, and this study was approved by the local ethics committee (CEUA - IBUSP / CAAE 65361317.2.0000.5464).

**Table 1.** Anthropometrics and other characteristics of the subjects.

| Subject | V<br>(Km/h) | HR<br>(bpm) | %<br>HRmax | Estimated maximum<br>HR (bpm) | Age<br>(years) | Height<br>(cm) | Weight<br>(Kg) | BMI<br>(Kg/m<sup>2</sup>) |
| --- | --- | --- | --- | --- | --- | --- | --- | --- |
| R.A. | 12.0 | 172 | 90 | 191 | 24 | 168 | 79 | 28.0 |
| F.C. | 12.0 | 150 | 84 | 178 | 43 | 163 | 71 | 26.7 |
| D.Q. | 11.0 | 167 | 87 | 193 | 22 | 166 | 53 | 19.2 |
| V.T. | 12.5 | 142 | 77 | 185 | 33 | 172 | 85 | 28.7 |
| C.S. | 11.0 | 166 | 88 | 188 | 29 | 174 | 84 | 27.7 |
| V.Y. | 11.0 | 155 | 83 | 186 | 31 | 185 | 98 | 28.6 |
| *Mean* | *11.6* | *159* | *85* | *187* | *30* | *171* | *78* | *26.5* |
| *s.d.* | *0.7* | *12* | *5* | *5* | *8* | *8* | *15* | *3.6* |

### Speed selection

A control velocity (V) was selected for each volunteer to be taken as a reference for calculating the other experimental velocities. V was chosen to elicit, at 0% incline, an HR of 77% to 95% of the maximum estimated HR of the volunteer. This range corresponds to “vigorous-intensity” according to the American College of Sports Medicine (GARBER et al., 2011). Maximum HR (HRmax) was estimated according to Tanaka et al. (2001) as HRmax = 208 - 0.7 × age in years

The range of speed and slope were selected following what runners usually encounter on the streets (that is, relatively low slopes) and such as the subjects would retain a running gait at the lowest velocity yet still be able to sustain the activity at the condition with the highest metabolic demand. Intermediate conditions were chosen according to pilot experiments to result in similar heart rates. We opted for an incremental protocol (that is, increasing speed) instead of random order or a decremental protocol, as suggested by Natali (2016) to be more adequate in studying HRV.

### Data acquisition

Subjects ran on a motorized treadmill at room temperature (average of 21.3 °C, s.d. = 2.3 °C) and relative humidity of 62% (s.d. = 8.7%). Experiments were performed indoors at the Department of Physiology, Bioscience Institute, University of São Paulo.

Data was acquired through Biopac Student Lab Pro MP36 (Biopac Systems, Inc.), at a sampling rate of 1000 Hz, using surface electrodes. The electrocardiographic signal was obtained with electrodes in a modified CM5 thoracic configuration. Equipment was set at AHA setup, with a low pass filter of 0.05 Hz and a high pass of 100 Hz.

### Experimental protocol

Volunteers were asked to avoid strenuous training and to abstain from alcohol and caffeine in the 24 hours preceding each experimental session. Sessions took place between 7 a.m. - 10 a.m., over 4 weeks or less.

Once electrodes were placed, the volunteer performed a 6-minute warm-up (3 minutes walking at 4 km/h followed by 3 minutes running at 0.8 V). The treadmill was then stopped and contingent adjustments in electrodes and/or cables were performed. After such a procedure, the experimental runs began.

Experimental runs were composed of 4 consecutive 3-minute runs, performed at increasing speeds of 0.7 V, 0.8 V, 0.9 V, and 1 V. The slopes, randomly selected each day, were 0%, and 6%. Only one slope was performed at each visit to the laboratory. In total, volunteers engaged in 8 combinations of speed and slope over 2 non-consecutive days.

### Signal processing

The Signal was processed and analyzed using MATLAB R2015b (MathWorks Inc.) and/or R 3.4.3 software (R Project). The interval between successive R waves in the electrocardiographic signal (RR interval) was obtained and was visually inspected for abnormal/ectopic beats, which were excluded from the analysis. Abnormal/ectopic beats comprise less than 1% of the total beats in each one of the experiments.

The last 15 seconds of each run were summarily discarded to avoid effects associated with the transition between conditions. The RR segment analyzed was composed of 256 RR intervals obtained from the last minutes under a given combination of speed and slope.

The HR was measured for each volunteer under a certain combination of speed and slope. To make sensible comparisons among the individuals, the HR of the pair {speed, slope} was normalized to the HR in the reference velocity (pair {1V, 0%}), and named as relhr.

HRV was estimated through the standard deviation (sdnn) and root mean square (rms) of successive RR intervals (GUIDELINES, 1996). The estimators of HRV obtained for each subject were divided by the HR measured in that condition (named sdt and rmst). This procedure allowed us to visualize the behavior of the HRV regardless of the HR itself.

### Statistical analysis

We were unable to obtain data from the run {0.7 V, 6%} for the volunteer C.S. due to excessive noise in the record. Also, two runs were not completed because the subjects reached exhaustion ({1 V, 6%} for V.Y. and C.S.) The presence of outliers (considered as points that deviated 3 or more interquartile intervals from the mean values) was verified and none were detected. The normality of HR, sdt, and rmst was accessed through the Shapiro-Wilk test and by visual inspection. The critical p-value was set at 0.05.

To verify if there were differences in the results, we performed two-way ANOVA with speed and slope as independent variables. If the ANOVA test yielded significant results, we then proceeded to a Student-Newman-Keuls (SNK) test to verify which conditions were different from each other.

## RESULTS

Results obtained for HR, sdnn, and rms are presented in Table 2 in absolute values, relative to the run {1 V, 0%} (relative) and divided by HR (sdt, rmst). Relative HR (relhr), sdt, and rmst are also graphically presented in Fig. 1 – 3.

**Table 2.**
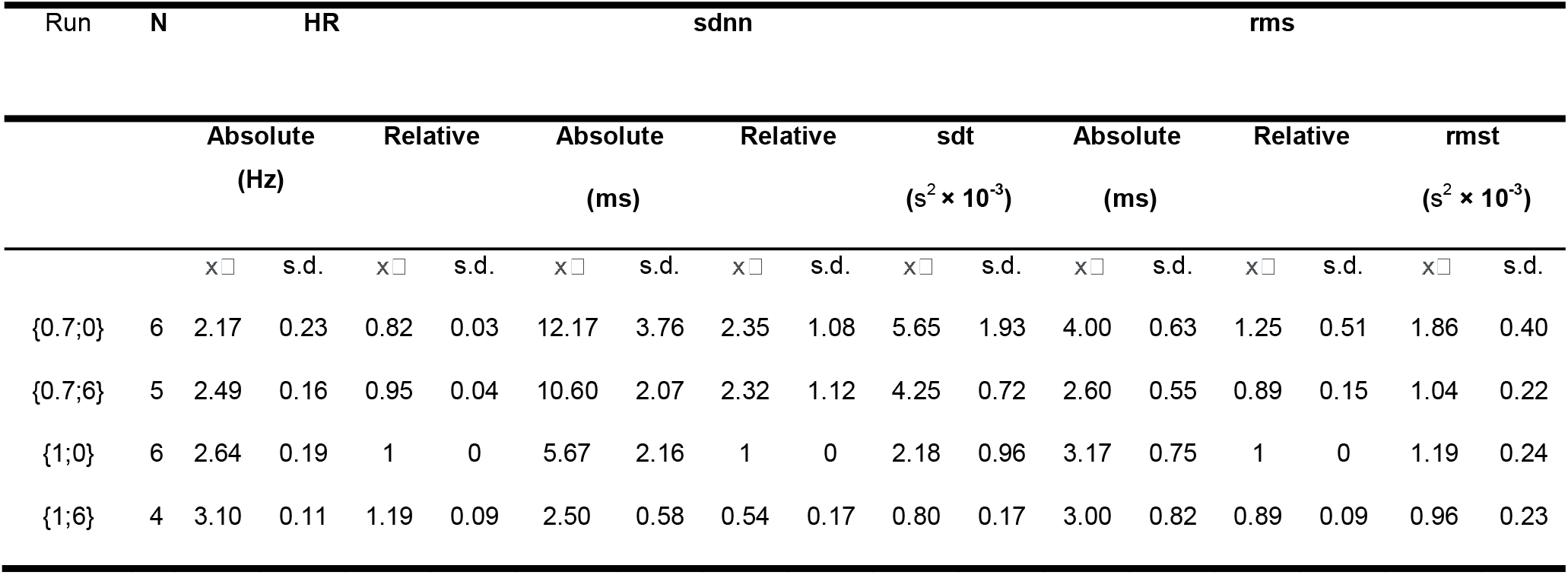
HR, sdnn and rms (mean - x□ - and s.d.) in absolute, relative to the run at {1 V, 0%} and divided by HR sdt, rmst) for the 4 experimental conditions. N indicates number of subjects analyzed under each condition.

**Fig. 1.**
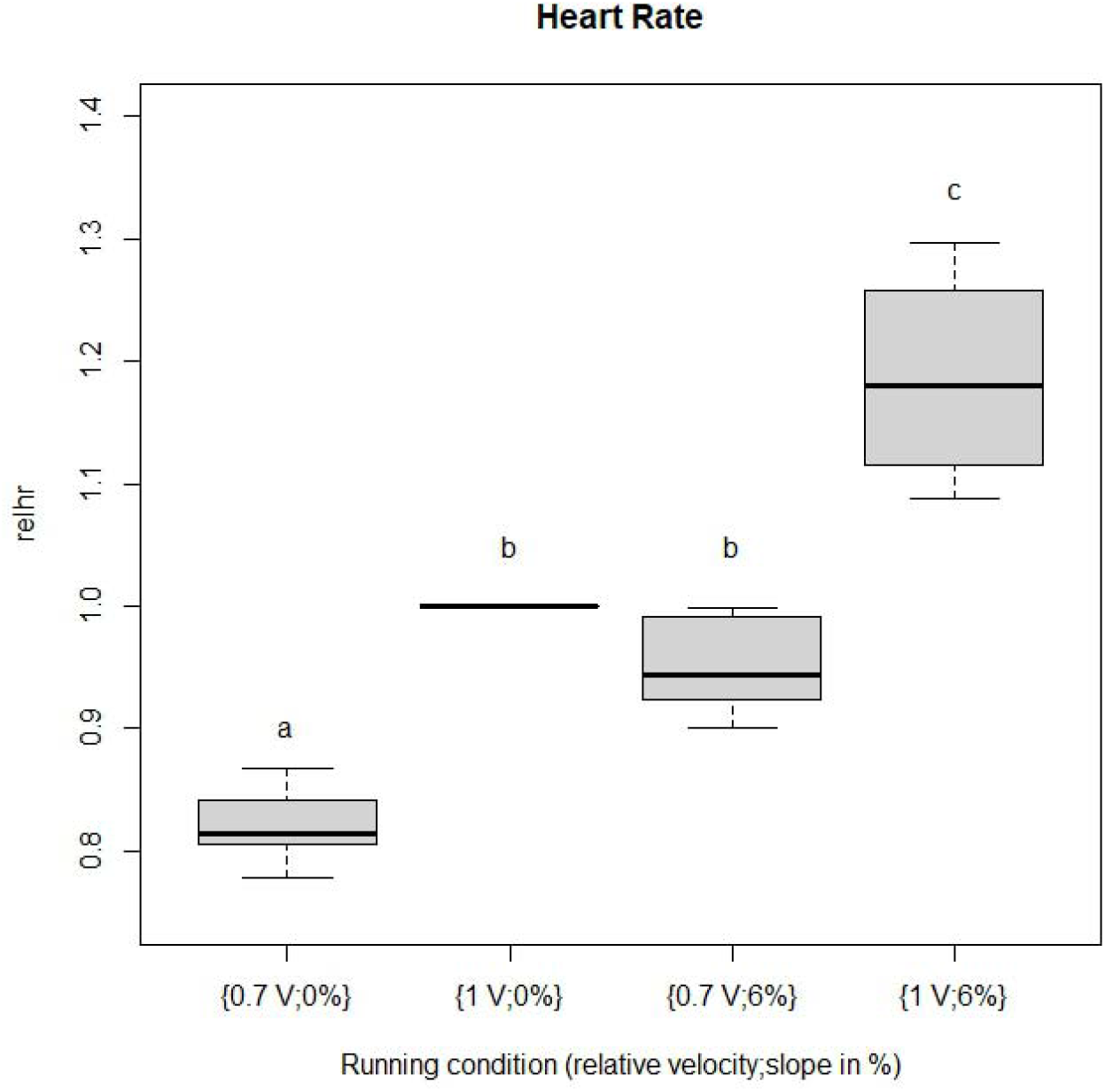
Relative HR. HR divided by the values observed in the run at 1 V speed, 0% slope (for {0.7 V, 0 %} and {1 V, 0%}, n = 6, for {0,7 V; 6%}, n = 5 and for {1 V; 6%}, n = 4). Different letters indicate which conditions are significantly different (or not) from the others according to the two-way ANOVA followed by SNK tests.

### relhr

Results from the ANOVA test indicated that, as expected, relhr responds to both speed (F = 50.30, p-value-< 0.0001) and slope (F = 99.20, p-value < 0.0001), and there is no significant interaction between the independent variables (F = 1.75, p-value = 0.20). SKN results (Fig. 1) show that the runs {0.7 V, 0%} and {1 V, 6%} are each different from the other 3 conditions, while the intermediate intensity runs, {0.7 V, 6%} and {1 V, 0%}, are no different from one another.

### sdt

Results from the ANOVA test indicated that sdt responds to both speed (F = 4.93, p-value = 0.04) and slope (F = 41.80, p-value < 0.0001), and there is no significant interaction between the independent variables (F = 0.00, p-value = 0.99). SNK results (Fig. 2) show that the runs performed at {0.7 V, 0%} and {0.7 V, 6%} are different from those at {1 V, 0%} and {1 V, 6%}.

**Fig. 2.**
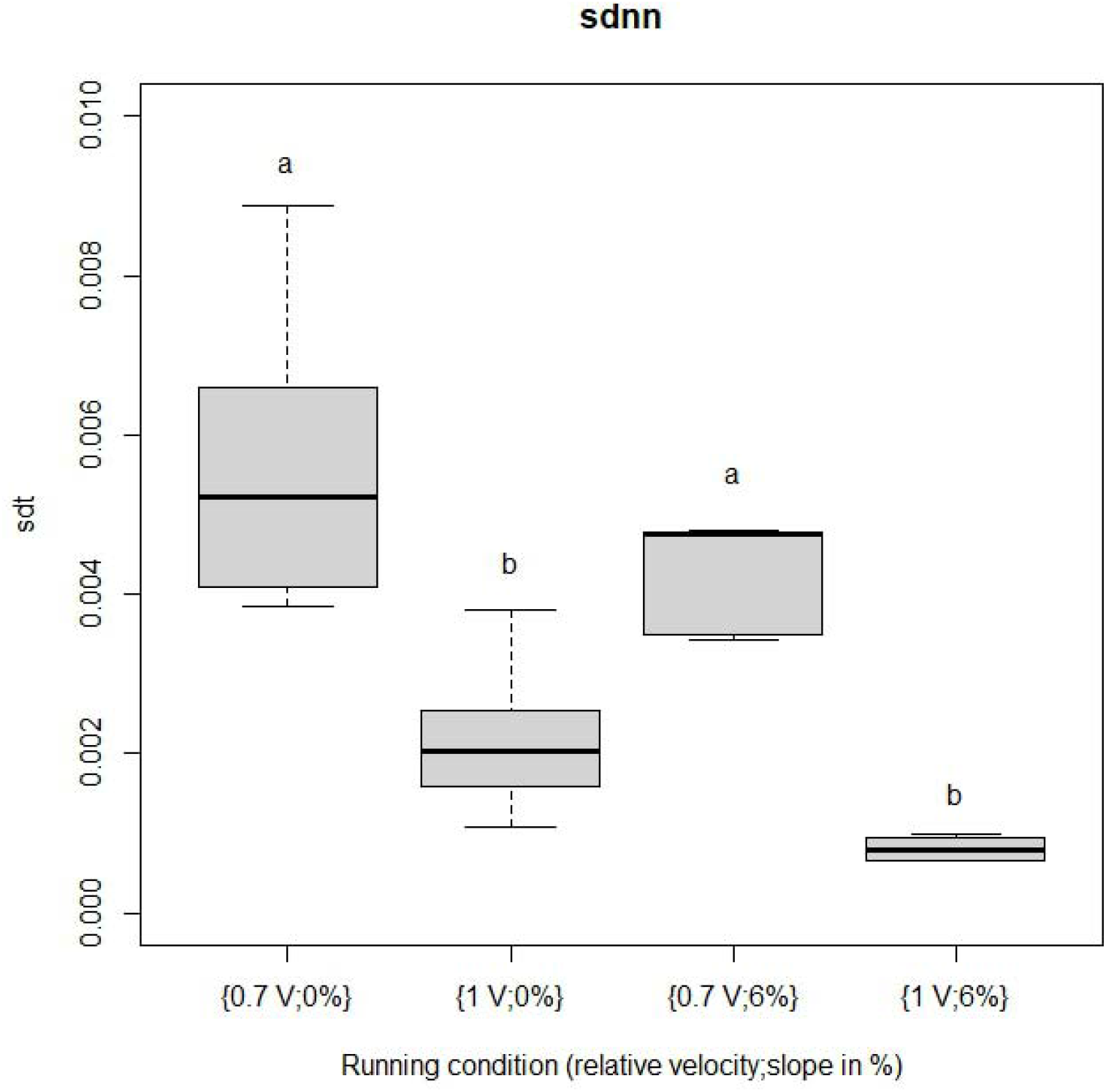
sdt (s^2^) for each run. sdt was calculated as sdnn (in seconds) divided by HR (Hz) (for {0.7 V, 0%} and {1 V, 0%} n = 6, for {0,7 V; 6%} n = 5 and for {1 V; 6 %}, n = 4). Different letters indicate which conditions are significantly different (or not) from the others according to the two-way ANOVA followed by SNK tests

### rmst

Results from the ANOVA test indicated that rmst responds to both speed (F = 16.36, p-value < 0.001) and slope (F = 10.91, p-value = 0.004), and there is significant interaction between the independent variables (F = 5.22, p-value = 0.036). SNK results (Fig. 3) show that the run {0.7 V, 0%} is different from the remaining 3 conditions, which, in turn, are no different from one another.

**Fig. 3.**
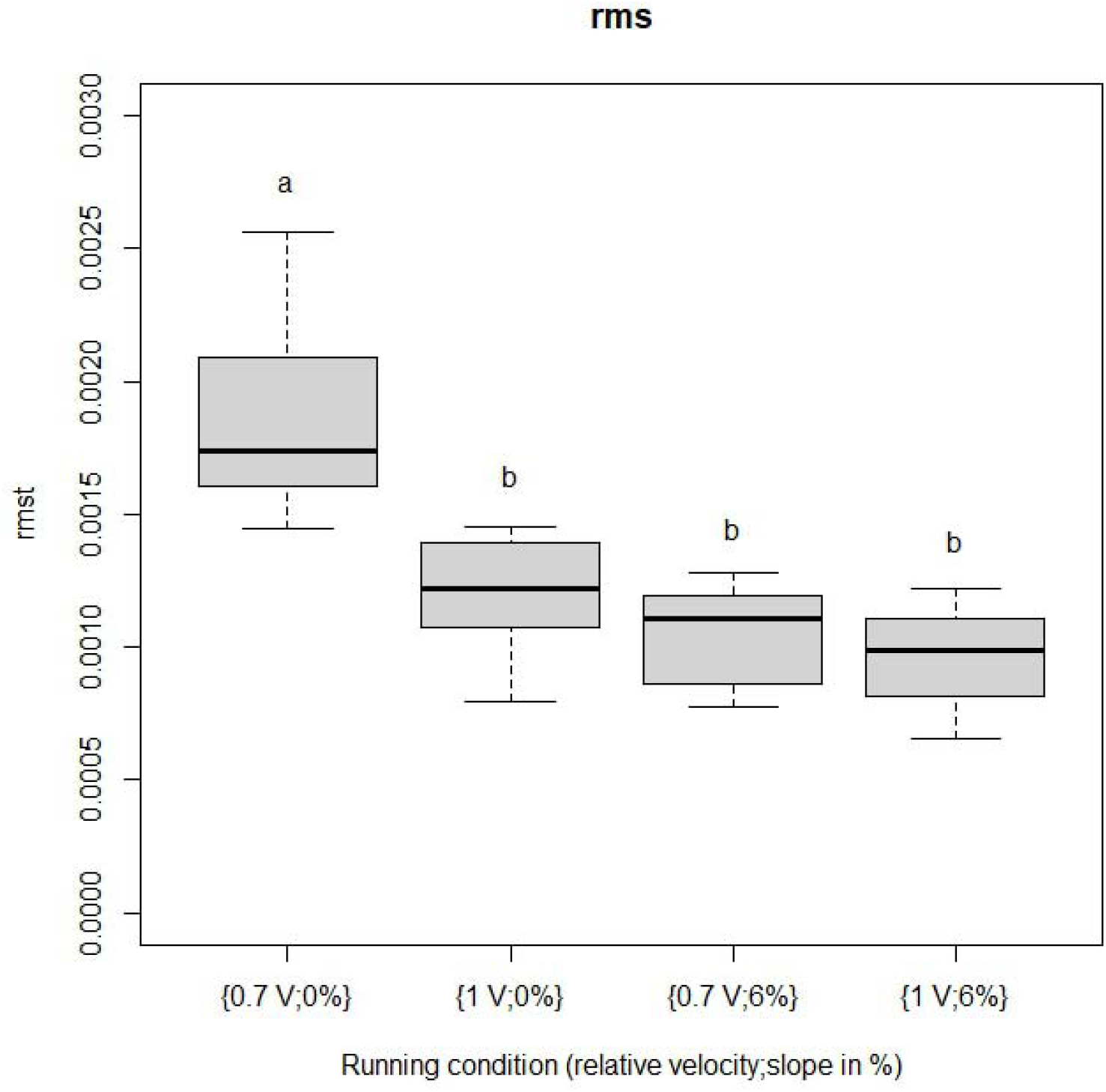
rmst (s^2^) for each run. rmst was calculated as rrms (in seconds) divided by HR (Hz) (for {0.7 V, 0%} and {1 V, 0%}, n = 6, for {0,7 V; 6%}, n = 5 and for {1 V; 6%}, n = 4). Different letters indicate which conditions are significantly different (or not) from the others according to the two-way ANOVA followed by SNK tests.

## DISCUSSION

As explained in the Introduction, heart rate variability is associated with heart rate itself, which, in turn, responds to metabolic demand irrespectively to the origin of such a demand. We aimed to test whether HRV would be sensitive to changes in slope or speed under a similar workload, i.e., under similar HR.

Even at the lowest intensity runs {0.7 V, 0%}, the subjects attained a mean HR of 130 BPM, which corresponds to approximately 70% of the estimated maximum HR for the group of volunteers. At such exercise intensity, the influence of the parasympathetic nervous system is likely restricted (Carter et al., 2003; Perini & Veicsteinas, 2003) and the sympathetic branch and hormonal adjustments are expected to play a more significant role in the control of HR. For instance, Kindermann et al. (1982) measured epinephrine and norepinephrine concentration following aerobic (mean HR of 176 BPM) and anaerobic (156% maximum exercise capacity – HR up to 185 BPM) activities (exercise intensities are approximately comparable with the lowest and highest intensity runs adopted in our study in terms of HR). They report a 15-fold increase in adrenaline and noradrenaline through the course of the aerobic activity, and a 4- to 9-fold increase in the anaerobic exercise. Therefore, we can assume that our protocol was effective in causing relevant parasympathetic withdrawal and an increase in sympathetic tone.

In our results, HR increased with both speed and slope, as expected, with no interaction between these variables. The intermediate conditions, that is, the combinations of maximum speed and minimum slope {1 V, 0%} and of minimum speed and maximum slope {0.7 V, 6%}, resulted in heart rates no different from one another (Fig. 1), allowing us to compare the sensitivity of cardiac variability facing different biomechanical loads.

HRV estimated as sdt and rmst, i.e., sdnn and rms estimators divided by the heart rate at the time of measurement, decreased with increasing speed or slope, with no interaction between these independent variables. rmst was higher on the lowest metabolic demand run (1.86 × 10-3 s^2^) and decreased by 44% (1.04 × 10-3 s^2^), 36% (1.19 × 10-3 s^2^), and 48% (0.96 × 10-3 s^2^) at the {0.7 V, 6%}, {1 V, 0%} and {1 V, 6%} runs, respectively (mean decrease of 43%). rmst for all values but the one obtained in the {0.7V, 0%} run are no different from one another (Fig 3). Thus, this estimator did not show a distinct sensitivity to variations in speed or slope.

sdt, on the other hand, was more sensitive to changes in speed than in slope. As shown in Figure 2, when relative velocity was increased from 0.7 to 1, there was a significant change in sdt, with values dropping nearly 61%, from 5.56 × 10-3 s^2^ to 2.18 × 10-3 s^2^. However, when the same HR was attained through a higher slope, sdt was 4.25 × 10-3 s^2^, not significantly different from that obtained at the {0.7 V, 0%} condition (Fig. 1). This finding opposes our hypothesis that HRV would decrease proportionally only to the HR and indicates that sdt is sensitive to other factors. The behavior of sdt may reflect the different stride patterns associated with changes in speed or slope, the ventilatory frequency, or a combination of those, since stride and ventilatory frequency are often coupled (SUMI et al., 2006).

Many studies reported the persistence of components associated with ventilatory frequency in HRV under all exercise intensities (Perini et al., 1990; Perini & Veicsteinas, 2003; Sumi et al., 2006). (Bailon et al. (2013) hypothesized the presence of estimators associated with ventilatory frequency under medium-high intensity exercise reflects the mechanical stretching of the sinus node due to exercise-induced hyperventilation above the anaerobic threshold. During inspiration, the reduction of intrathoracic pressure would lead to a greater venous return and, consequently, a greater atrial filling. This, in turn, leads to stretching of the SA node, raising HR. Therefore, the modulatory effect of ventilatory frequency on HRV during exercise would be mainly associated with a mechanical, rather than a neural, effect.

A cadence component is also relevant in modulating HR, as previously demonstrated, either through HRV analysis or more usually as synchronization between the step cycle and occurrence of the R wave (Alikhani et al., 2017; Blain et al., 2009; Kirby et al., 1989). One possible explanation to cardiolocomotor synchronization is of a mechanical origin, in which rhythmic muscle contraction modulates venous return (BLAIN et al., 2009). Under dynamic muscle contractions, venous return presents a pulsatile pattern, increasing during contraction and decreasing in the relaxation phase (Folkow et al., 1970). Such a hypothesis is further supported by the proportional relationship observed between power in the cadence component and workload (Blain et al., 2009). Additionally, as reviewed by Fisher (2014), the activation of mechanoreceptors on skeletal muscles leads to an increase in HR through suppression of parasympathetic activity. Metaboreceptors in muscular tissue evoke an increase in HR as well but via activation of the sympathetic branch. Given the exercise intensity in our experiments, the latter mechanism likely plays a more significant role in the HRV changes observed.

In summary, our experiments show that HRV decreases with both speed and slope, even when estimators were relative to the heart rate itself. On the one hand, the relative rms was not sensitive to the running conditions, reaching similar values to all but the lowest intensity run. On the other hand, relative sdnn decreased when speed was increased, but not when the same HR was attained through increments in slope. Therefore, we can conclude that HRV responds differently to increments in speed or slope during running, despite similar HR. This conclusion is relevant from a biological perspective, and might also play a role when considering the training routine of runners in general.

## CONCLUSION

Our results strongly suggest that cardiac control operates in a sensitive manner in the face not only of metabolic demand but also due to the mode of the changes imposed on such a demand. As far as we know, this is the first time this phenomenon is clearly evidenced.

## ACKNOLEDGEMENTS

We are grateful for discussions and insights given by Eric C. Becman, and Paulo N. Starzynski regarding the experimental protocol and analysis.

